# Autophagic cargo in Lewy bodies: are Lewy bodies a compartment for spatial protein quality control?

**DOI:** 10.1101/2023.09.24.559164

**Authors:** Phoebe Conod, Nicola Charlesworth, Pawel Palmowski, Andrew Porter, Lauren Walker, Omar El-Agnaf, Viktor Korolchuk, Tiago F. Outeiro, Daniel Erskine

## Abstract

Lewy bodies are neuropathologically associated with Lewy body dementia (LBD), but little is known about why they form or their role in the disease process. We previously reported Lewy bodies are a common feature of older individuals with primary mitochondrial diseases. However, as they are not an invariant finding, understanding differences between those with and without Lewy bodies may provide insights into factors that govern the formation of Lewy bodies in Lewy body disease (LBD). The present study sought to investigate whether deficient mitophagy in the context of mitochondrial dysfunction may underlie Lewy body formation. *Post-mortem* tissue was obtained from the cingulate gyrus and dorsal motor nucleus of the vagal nerve (DMV) of mitochondrial disease cases with Lewy bodies, primary mitochondrial disease cases without Lewy bodies, and control cases, in addition to LBD cases as comparison. An array of mitophagy and autophagy markers were quantified in 50 individual neurons per cingulate gyrus and all neurons per DMV using immunofluorescent analysis. No significant differences were found between groups, although there was a striking enrichment of markers of autophagic mitochondria and autophagic vesicles within Lewy bodies. Evaluation of diffuse α-synuclein aggregates, thought to precede Lewy body formation, suggested only autophagic mitochondria were present in early aggregates, perhaps suggesting sequestration of dysfunctional mitochondria is an early step in Lewy body formation. To characterise the composition of Lewy bodies, discovery proteomics was performed on isolated insoluble proteins from frozen cingulate gyrus, which identified up-regulation of markers of aggresomes, a regulated cellular response that occurs when protein degradative pathways become overwhelmed, a mechanism of spatial protein quality control (sPQC). Taken together, these findings are consistent with impairment of cellular waste handling pathways in Lewy body-bearing neurons, and that the formation of a Lewy body could be a deliberate cellular response to compartmentalise such waste.

## INTRODUCTION

Lewy body disease (LBD) is an umbrella term for Parkinson’s disease (PD), Parkinson’s disease dementia (PDD) and dementia with Lewy bodies (DLB), a continuum of clinically and pathologically related entities that are thought to collectively comprise the second most common form of neurodegenerative disorder after Alzheimer’s disease [1]. LBD are differentiated based on the temporal sequence in which clinical features manifest, PD presents with motor symptoms that can become a cognitive disorder (PDD), whilst DLB patients initially present with cognitive symptoms and movement disorders can occur later [2]. LBDs are incurable and treatment options are limited to symptomatic therapies of limited benefit to patients.

Although LBDs are clinically distinguishable from one another, they share a unifying neuropathological feature in the presence of intracellular accumulations of the protein α-synuclein in neurons, termed Lewy bodies [3]. The association between LBD and the presence of Lewy body pathology has led to a prevailing view that Lewy bodies have a pathogenic role [4], and the process of α-synuclein aggregation into Lewy bodies has become a major therapeutic target in LBD [5]. Nevertheless, the evidence linking Lewy bodies and the neurodegeneration that is thought to be critical in LBD is limited, with previous studies reporting neurodegeneration precedes Lewy body formation in critical brain regions [6], the presence of neurodegeneration in regions without Lewy bodies and preservation in regions with Lewy bodies [7], and the absence of a relationship between the abundance of Lewy bodies and neuronal loss in the substantia nigra [8]. Therefore, there is compelling evidence from *post-mortem* neuropathology studies that the association between Lewy body formation and cell death in LBD is, at best, not straightforward.

Although most neuropathological studies investigating the role of Lewy bodies have compared the density of Lewy bodies with degree of neuronal loss, an increasing number of studies have directly compared neurons with Lewy bodies to proximal neurons without Lewy bodies. Although almost all of these studies have examined markers of components of the mitochondrial respiratory chain, they have showed a consistent picture of deficiencies in Complex I of the mitochondrial respiratory chain in neurons without Lewy bodies, and apparent preservation of Complex I in Lewy body-bearing neurons [9-11]. Our study of the mitochondrial respiratory chain in the nucleus basalis of Meynert of LBD cases identified higher expression of Complex I in Lewy body-bearing neurons but also generally higher mitochondrial mass in LBD cases [11]. A contemporaneous study using advanced imaging further reported that dystrophic, and thus likely dysfunctional, mitochondria were observed within Lewy bodies [12], led us to speculate that Lewy bodies may play a protective role in sequestering damaged mitochondria in the context of deficient mitochondrial autophagy (mitophagy) [13].

Although mitochondrial dysfunction has been long implicated in the pathogenesis of LBD, how it could contribute to Lewy body formation is not yet clear. Mitochondrial dysfunction generally up-regulates the cellular catabolic process of autophagy, likely to increase mitochondrial biogenesis and the pool of healthy mitochondria [14]. However, whether enhanced autophagy can be sustained long-term is unknown. We have previously reported Lewy body pathology to be a common feature of individuals with primary mitochondrial diseases, but it is notable that Lewy bodies were only found in individuals with mitochondrial disease in advanced age (over 50 years old) [15]. Ageing is known to lead to impairment of various aspects of the autophagic machinery, including lysosomal function and ATG proteins [16]. Therefore, one could hypothesise that mitochondrial dysfunction on its own is not sufficient to lead to Lewy body pathology, but rather it is the combination of mitochondrial dysfunction and impaired mitophagy. Such a situation would lead to the accumulation of damaged mitochondria, which are harmful to neuronal health by producing reactive oxygen species, and thus the formation of a Lewy body to encapsulate such waste products in an insoluble structure may be a protective mechanism.

The present study sought to interrogate the hypothesis that mitophagy is perturbed in mitochondrial disease cases with Lewy bodies, and the formation of Lewy bodies may serve to encapsulate damaged mitochondria, a cellular mechanism of spatial protein quality control (sPQC) [17]. The aim of this study was to quantify markers of mitophagy and autophagy in individual neurons of mitochondrial disease cases and controls, and directly compare neurons with and without Lewy bodies.

## METHODS

### Case selection

Human brain tissue was obtained from the Newcastle Brain Tissue Resource and was chosen on the basis of availability of tissue. Formalin fixed paraffin-embedded (FFPE) and frozen tissue was obtained from the cingulate gyrus, chosen due to its predilection to Lewy body pathology, from control cases (N=8), mitochondrial disease cases with Lewy bodies (N=4), mitochondrial disease cases without Lewy bodies (N=3) and PDD cases for comparison (N=2). When available, FFPE tissue was obtained from the corresponding dorsal motor nucleus of the vagal nerve (DMV) from the same cohort of cases, though exhaustion of tissue in some cases reduced the cohort size (control, N=7; mitochondrial disease cases without Lewy bodies, N= 2; mitochondrial disease cases with Lewy bodies, N=4; PDD cases, N=2). Cohort details are contained in Table 1. MitoLB 3 was initially diagnosed with CPEO of mitochondrial aetiology *intra vitam* due to a heterozygous variant in TWNK; however, subsequent investigations *post-mortem* also indicated over 11 repeats in PABPN1, consistent with a diagnosis of oculopharyngeal muscular dystrophy (OPMD), which could contribute to CPEO symptoms. Nevertheless, as a mitochondrial aetiology is long-established in OPMD, we have continued to include this case amongst mitochondrial disease cases in this study [18, 19].

**Table 1:**
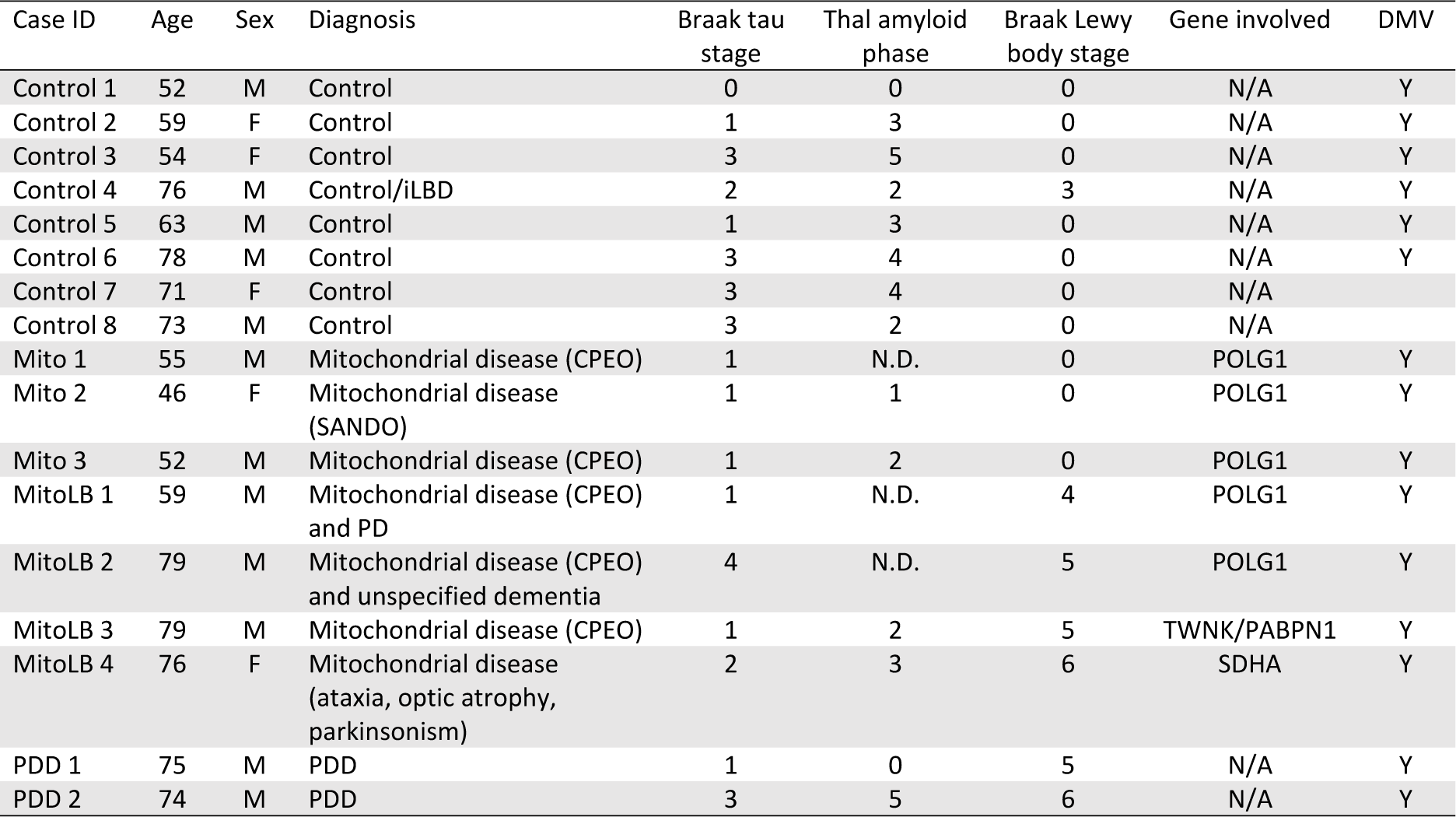
Demographic information on the cohort used in the present study. Braak tau stage, Thal amyloid phase and Braak Lewy body stage are based on previously reported staging systems [43-45]. “iLBD” refers to incidental Lewy body disease, “CPEO” to chronic progressive external ophthalmoplegia, “SANDO” to sensory ataxic neuropathy, dysarthria, and ophthalmoparesis, “PDD” to Parkinson’s disease dementia, “N.D.” to not determined. “DMV” indicates whether the dorsal motor nucleus of the vagal nerve was available for this case or not.

### Immunofluorescent staining of tissues

FFPE sections from the cingulate gyrus and DMV were cut at 5µm thickness for immunofluorescent analysis. Sections were heated in a 60°C oven for 20 minutes before dewaxing in two changes of HistoClear for 15 minutes each. Sections were then rehydrated through a graded series of ethanol solutions until water. Antigen retrieval was optimised for each antibody array individually but was primarily performed by boiling sections in citrate buffer pH6 for 10 minutes, with the addition of 5 minutes in formic acid afterwards in some cases as noted in Table 2. Sections were blocked in 10% normal goat serum in tris-buffered saline with Tween 20 (TBST) for one hour at room temperature overnight before application of primary antibody solutions as specified in Table 2 and incubated overnight at 4°C. Primary antibodies were washed off on the second day and sections were incubated in secondary antibody solutions for one hour at room temperature Table 2 prior to counter-staining in 3% Sudan black B solution to attenuate autofluorescence and mounting in ProLong Gold Antifade mountant. To ensure inter-assay variation did not confound results, all slides to be compared were stained at the same time, and staining was compared to sections stained only with secondary antibodies to ensure fluorescent signals were the result of primary antibody staining.

**Table 2:** List of antibodies used.

| Antibody target | Manufacturer/catalogue number | Host species | Isotype | Dilution | Antigen retrieval |
| --- | --- | --- | --- | --- | --- |
| **Primary antibodies** |  |  |  |  |  |
| NeuN | Abcam ab104224 | Mouse | IgG2b | 1:200 | Citrate pH6 |
| NeuN | Abcam ab279296 | Mouse | IgG2a | 1:500 | Citrate pH6 |
| Aggregated α-synuclein | Omar El-Agnaf syn-O2 | Mouse | IgG1 | 1:1000 | Citrate pH6 |
| Aggregated α-synuclein | Omar El-Agnaf syn-F2 | Mouse | IgG2a | 1:1000 | Citrate pH6 |
| α-synuclein phosphorylated at serine 129 | Abcam ab51253 | Rabbit | IgG | 1:500 | Citrate pH6 |
| Ubiquitin phosphorylated at serine 65 | Merck ABS1513-I | Rabbit | IgG | 1:100 | Citrate pH6 |
| BNIP3 | Abcam ab109362 | Rabbit | IgG | 1:500 | Citrate pH6 |
| VDAC1/porin | Abcam ab14734 | Mouse | IgG2b | 1:100 | Citrate pH6 |
| 3-nitrotyrosine | Abcam ab61392 | Mouse | IgG2a | 1:2000 | Citrate pH6 |
| ATG5 | Abcam ab238092 | Mouse | IgG1 | 1:25 | Citrate pH6 |
| LC3B | Abcam ab48394 | Rabbit | IgG | 1:500 | Citrate pH6 |
| HDAC6 phosphorylated at serine 22 | GeneTex GTX55403 | Rabbit | IgG | 1:100 | Citrate pH6 + formic acid |
| Gamma tubulin | Thermo MA1-850 | Mouse | IgG2b | 1:200 | Citrate pH6 + formic acid |
| **Secondary antibodies** |  |  |  |  |  |
| Anti-mouse IgG1 – AF647 | Thermo A21240 | Goat |  | 1:100 |  |
| Anti-mouse IgG2a – AF546 | Thermo A21133 | Goat |  | 1:100 |  |
| Anti-mouse IgG2b – AF488 | Thermo A21141 | Goat |  | 1:100 |  |
| Anti-rabbit IgG – AF405 Plus | Thermo A48254 | Goat |  | 1:100 |  |

### Analysis of abundance of mitophagy/autophagy markers

To measure the abundance of individual markers, analysis was performed of individual neurons in the cingulate gyrus and DMV. Sections were imaged on a Leica SP8 confocal microscope, with images captured of groups of neurons at 63x magnification for subsequent analysis. Images were then imported into FIJI software and stacks made of the four channels imaged, enabling a region of interest to be drawn in one channel and analysed in another. For the present analysis, regions of interest were determined based on NeuN staining, or the combination of NeuN and α-synuclein to identify intracellular Lewy bodies. NeuN images had a region of interest drawn around the cytoplasm and excluded the nucleus, if visible, to ensure only cytoplasmic abundance was measured (Figure 1). The intensity of the other channels was then measured in the annotated region of interest to measure the various markers within the cytoplasm of the neuron in question. For analysis of the abundance of the various mitophagy/autophagy markers with neurons for comparison across groups, we aimed to analyse 50 neurons per cingulate gyrus per case, and every neuron per DMV per case.

**Figure 1:**
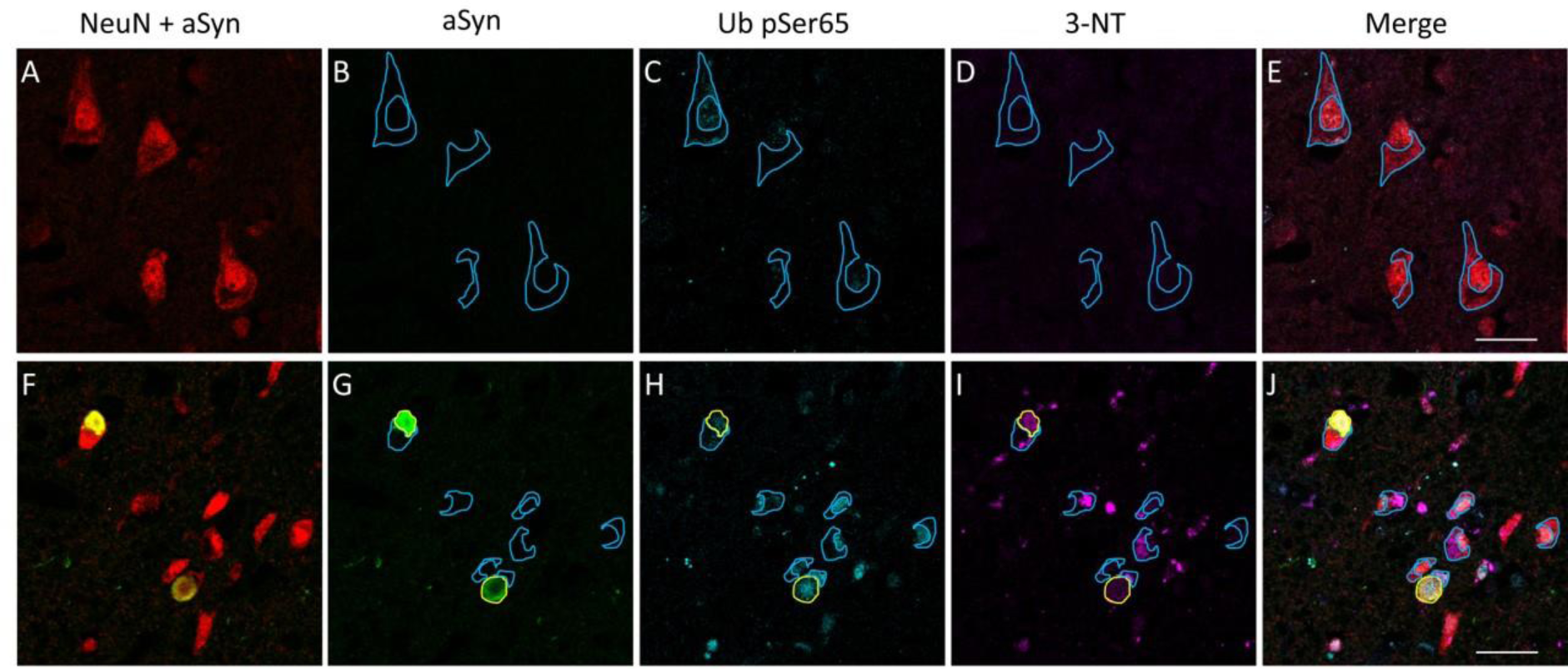
Methods of analysis in histological sections. A region of interest was drawn around the cytoplasm of neurons labelled by NeuN, excluding the nucleus if it was visible, to determine the abundance of a given marker within individual neurons (blue lines). α-synuclein accumulations were similarly delineated (yellow lines), thus enabling comparisons of neurons, Lewy body-bearing neurons and Lewy bodies themselves. Mean intensity measured were employed that normalised size of region of interest, to ensure different sizes of structures did not influence results. Scale bars = 30 µm.

To determine differences in neurons with Lewy bodies, or within Lewy bodies themselves, images were also obtained of neurons with α-synuclein aggregates. In this instance, every neuron with an α-synuclein aggregate was captured within a given slide of a particular case and, although we would have liked to separate aggregates on the basis of morphology, to ensure adequate statistical power we characterised all spherical intensely-stained deposits as “Lewy bodies” and all unstructured and diffuse deposits as “diffuse aggregates”, as we have done previously [13]. As with neurons without inclusion bodies, only the cytoplasm was measured, omitting both the nucleus and the Lewy body, and with the Lewy body analysed on its own (Figure 1).

Data generated for each cell type or structure per case were log-transformed to improve data normality, and z scores generated for cases based on the control distribution. Although many previous studies have pooled neurons on the basis of disease group, we elected not to do this as it would violate the principle of the independence of observations that underlies many statistical tests. Therefore, we generated mean values per case for group-level analysis, and also compared individual cases to the control distribution.

### Mass spectrometry analysis of insoluble proteome

To determine the composition of Lewy bodies in cases in the present study, insoluble proteins were isolated using centrifugation and subject to data independent acquisition mass spectrometry. Frozen tissue analysis was only performed on cingulate gyrus samples due to abundance of tissue necessary for the isolation procedure. To isolate insoluble proteins, brain samples were first homogenised in ten volumes 0.2M triethylammonium bicarbonate buffer using a rotator-stator homogeniser. Samples were then centrifuged at 16,000 xg for 30 minutes at 4°C and the supernatant removed, prior to three washes in Tris-HCl solution and resuspension of the pellet in Tris-HCl with 5% SDS. Samples were centrifuged for 30 minutes at 16,000 xg and washed three times in 2% SDS, prior to resuspension in XX sarkosyl for overnight incubation at 4°C. Samples were centrifuged at 16,000 xg and the pellet washed and solubilised in 5% SDS using a PreCelLys bead beater system. The solubilised pellet was then subject to mass spectrometry-based analysis, using data independent acquisition.

Samples were analysed by LC−MS/MS using an Ultimate 3000 Rapid Separation LC (RSLC) nano LC system (Thermo Corporation) coupled with an Exploris 480 Quadrupole-Orbitrap Mass Spectrometer (Thermo Fisher Scientific, Waltham, MA, U.S.A.). 1ug of peptide mixture was loaded, first onto an Acclaim PepMap100 C18 LC Column (5 mm Å∼ 0.3 mm i.d., 5 μm, 100 Å, Thermo Fisher Scientific) at a flow rate of 10 μL min−1 maintained at 45 °C and then then separated on 50 cm RP-C18 µPAC™ column (PharmaFluidics), using a 60min gradient from 97 % A (0.1% FA in 3% DMSO) and 3% B (0.1%

FA in 80% ACN 3% DMSO), to 35 % B, at a flow rate of 400 nL min−1. The separated peptides were then injected into the mass spectrometer via Thermo Scientific μPAC compatible EasySpray emitter at the Ion Transfer Tube temperature of 320oC, spray voltage 1500V and analysed using data independent (DIA) acquisition. The total LCMS run time was 90min. For the full scan mode, the MS resolution was set to 60000, with a normalized automatic gain control (AGC) target of 300%, Dynamic maximum injection time and scan range of 390−1600 m/z. DIA MSMS were acquired with 49, variable size windows covering 390-1621 m/z range, at 30000 resolution, AGC target set to 3000%, dynamic maximum injection time and normalized collision energy level of 30%.

The acquired data has been analysed in DIA-NN version 1.8 [20] against human proteome database (Uniprot 3AUP00000564-2022.10.20) combined with common Repository of Adventitious Proteins (cRAP), Fragment m/z: 200-1800, enzyme: Trypsin, allowed missed-cleavages: 2, peptide length: 7-52, precursor m/z 300-1800, precursor charge: 2 -5, Fixed modifications: carbamidomethylation(C), Variable modifications:Oxidation(M),Acetylation(N-term). The results were then exported as a text file, reformatted using in house written R scripts and further processed in The Perseus software platform version 1.6.15.0 [21].

### Determination of presence of HDAC6 pS22 and γ-tubulin in Lewy bodies

To determine whether Lewy bodies and diffuse α-synuclein aggregates incorporate the aggresome markrs, HDAC6 pS22 and γ-tubulin, sections stained with these antibodies in addition to NeuN and α-synuclein were imaged and assessed by two raters (PC and DE). Immunopositivity for either was determined on the basis of their presence within the area occupied by the Lewy body or diffuse aggregate, in comparison to the cells in which they were located and presented as percentages of the total. This analysis was performed in this way to dichotomise the presence or absence of these markers within Lewy bodies, as continuous data would not have addressed the question of the proportion of aggregates that manifest these markers.

## RESULTS

### Demographics

Mitochondrial disease patients were unavoidably younger than any available cases with Lewy body pathology; however, to ensure age had no impact on any results we included both young and old controls. Comparison of young and old controls demonstrated no significant difference on any variable (Supplementary Figure 1), so they were combined for subsequent analyses. Sex was not significantly different across groups (χ²=0.744, p=0.689).

### Mitophagy markers are selectively incorporated into Lewy bodies

Evaluation of expression levels of the mitophagy markers pUbSer65 and BNIP3, the nitrosative stress marker 3-nitrotyrosine, and the general mitochondrial marker VDAC1/porin in DMV and cingulate gyrus neurons demonstrated no differences at the group level (Supplementary Figure 2). Comparison of neurons with and without Lewy bodies identified a consistent picture of incorporation of the mitophagy markers BNIP3, pUbSer65 and 3-nitrotyrosine into Lewy bodies in both the DMV and cingulate gyrus (Figures 2-3). In contrast, the general mitochondrial marker VDAC1/porin was generally lower in Lewy bodies compared to neurons (Figure 3). In contrast, there were relatively few differences in the cytoplasmic expression of these markers in normal neurons compared to neurons harbouring Lewy bodies, though pUbSer65 was increased in cingulate Lewy body-bearing neurons and 3-nitrotyrosine was increased in Lewy body-bearing neurons in both DMV and cingulate gyrus (Figures 2 - 3). Taken together, these findings indicate selective incorporation of mitophagy markers, but not general incorporation of mitochondria, into Lewy bodies.

**Figure 2:**
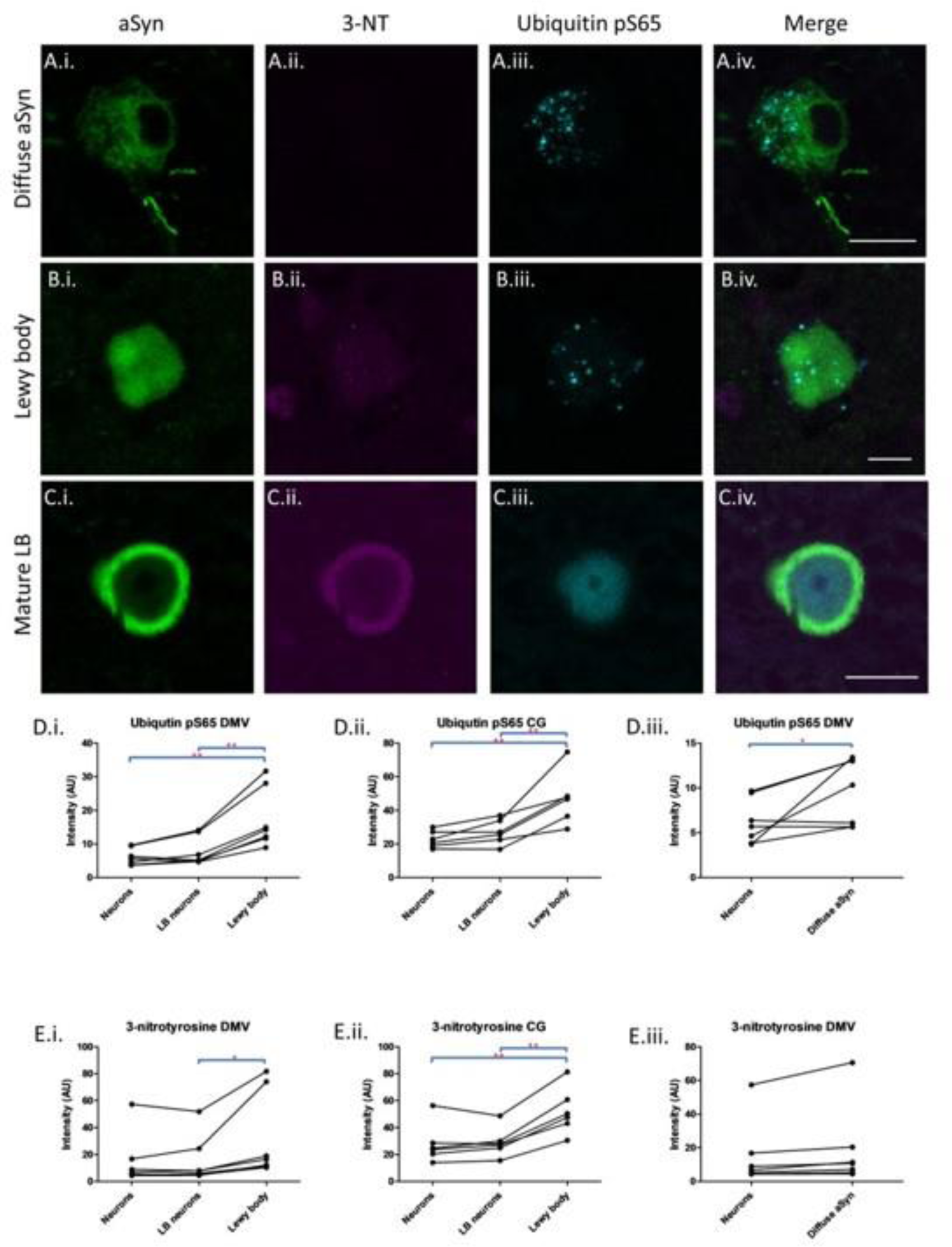
3-nitrotyrosine and ubiquitin pS65 in Lewy bodies. Representative images of diffuse α-synuclein aggregates (A.i.-A.iv.), Lewy bodies (B.i.-B.iv.) and mature O-shaped Lewy bodies (C.i.-C.iv.). 3-nitrotyrosine demonstrated diffuse staining within Lewy bodies (B.ii. and C.ii.) whilst ubiquitin pS65 labelled small structures that were thought to be autophagic mitochondria (A.iii., B.iii. and C.iii.). Quantitative analysis of abundance is shown in D.i. to E.iii., with each point representing the mean value per case and lines joining points from the same case to the trend of abundance across different cell types/structures within cases. Quantitative analysis of ubiquitin pS65 within LBD cases identified significantly higher abundance in Lewy bodies, Lewy body-bearing neurons and diffuse α-synuclein aggregates compared to neurons without Lewy bodies (D.i.-D.iii.). In contrast, 3-nitrotyrosine was more abundant in Lewy bodies compared to Lewy body-bearing neurons, but only higher compared to unaffected neurons in the cingulate gyrus and not significantly enriched in diffuse α-synuclein aggregates (E.i.-E.iii.). *p<0.05, **p<0.01.

**Figure 3:**
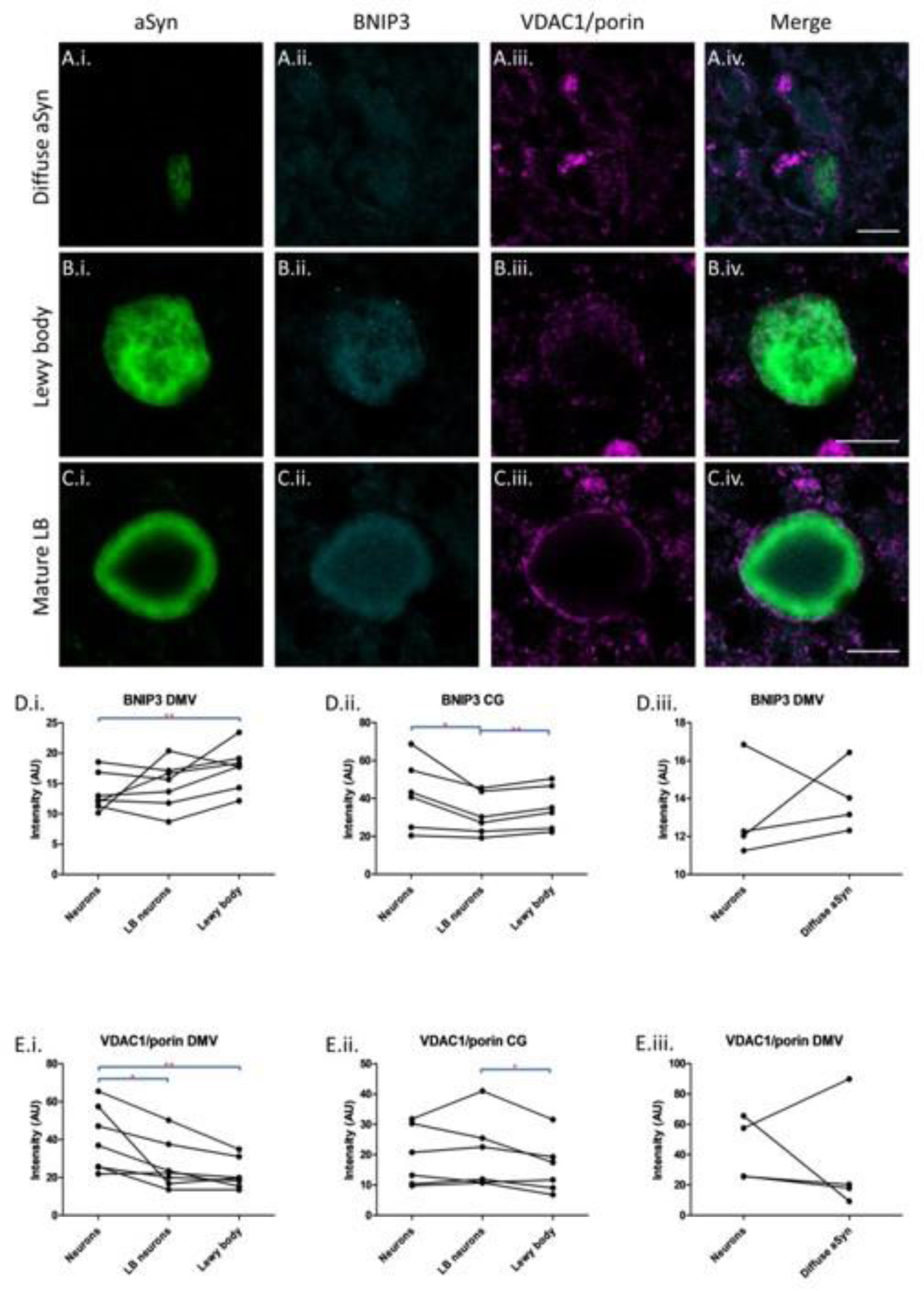
BNIP3 and VDAC1/porin in Lewy bodies. Representative images of diffuse α-synuclein aggregates (A.i.-A.iv.), Lewy bodies (B.i.-B.iv.) and mature O-shaped Lewy bodies (C.i.-C.iv.). Although BNIP3 was often present within some Lewy bodies, VDAC1/porin showed minimal co-localisation with all α-synuclein aggregates and a conspicuous absence from mature Lewy bodies, where it was often present as a ring around the Lewy body C.i.-C.iii. Quantitative analysis of BNIP3 within LBD cases demonstrated it was enriched in DMV Lewy bodies compared to unaffected neurons (D.i.); whereas, in the cingulate gyrus, BNIP3 was significantly reduced in the cytoplasm of Lewy body-bearing neurons compared to that of unaffected neurons and Lewy bodies themselves (D.ii.). BNIP3 also showed no consistent enrichment within diffuse α-synuclein aggregates (D.iii.). VDAC1/porin showed lower abundance in Lewy bodies compared to the cytoplasm of both unaffected neurons and Lewy body-bearing neurons from the same case in the DMV (E.i.) and lower abundance in Lewy bodies compared to Lewy body-bearing neurons within the cingulate gyrus (E.ii). VDAC1/porin was not consistently enriched in diffuse α-synuclein aggregates (E.iii.). *p<0.05, **p<0.01.

### Phagophores and autophagosomes are components of Lewy bodies

To determine whether incorporation of autophagic mitochondria into Lewy bodies is selective for mitochondria or extends to other autophagic cargo, we next studied the expression of the phagophore marker ATG5 and the autophagosome marker LC3B. As noted for mitophagy markers, no difference in abundance of either marker was noted at the group level (Supplementary Figure 2). However, when neurons with and without Lewy bodies were compared, we noted that both ATG5 and LC3B were enriched within Lewy bodies (Figure 4). It is also notable that in cingulate gyrus, we noted lower levels of both markers within Lewy body-bearing neurons which, combined with their enrichment in Lewy bodies, could lead one to speculate that they are being translocated from the cytoplasm to the Lewy body (Figure 4).

**Figure 4:**
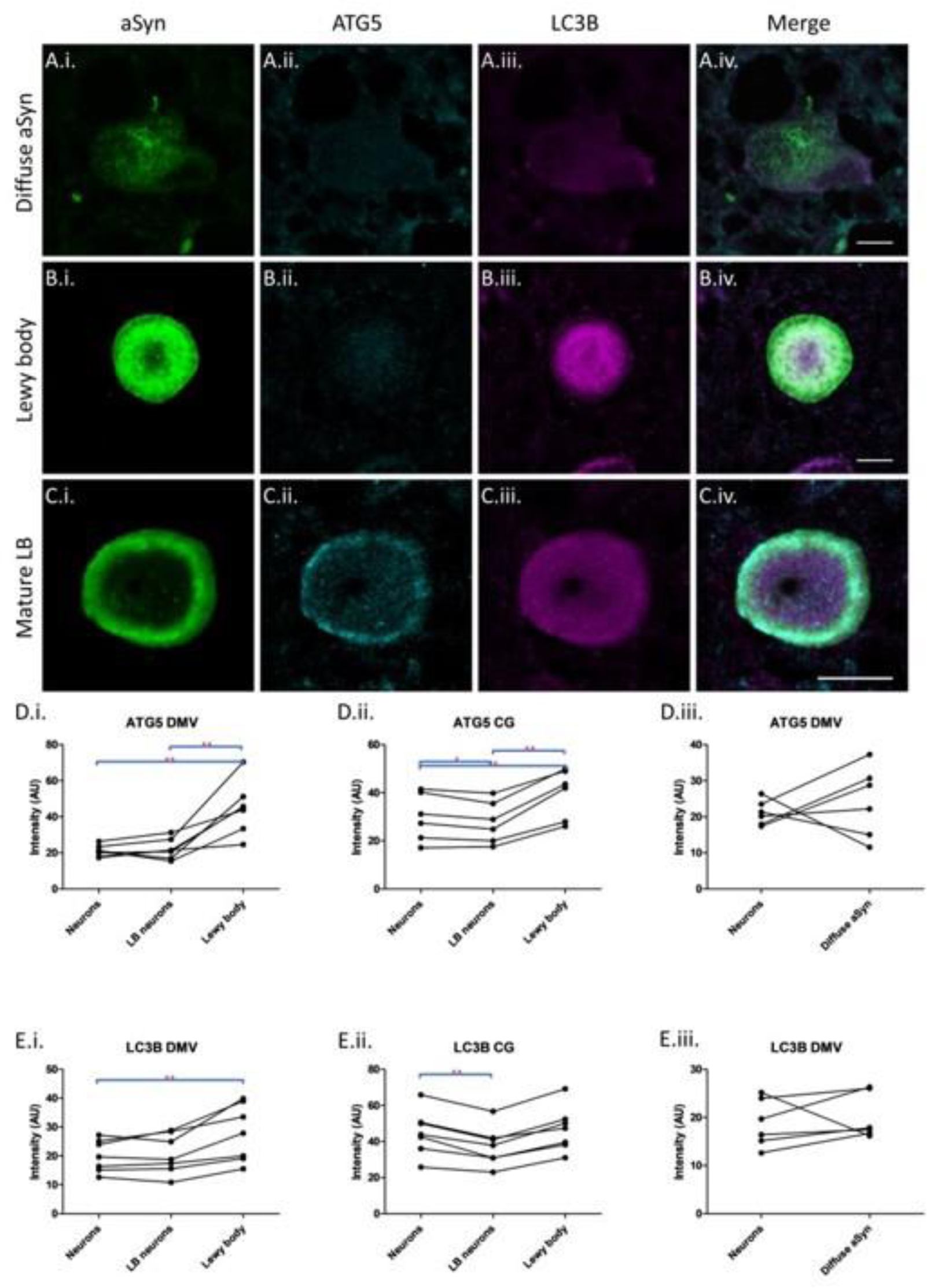
ATG5 and LC3B in Lewy bodies. Representative images of diffuse α-synuclein aggregates (A.i.-A.iv.), Lewy bodies (B.i.-B.iv.) and mature O-shaped Lewy bodies (C.i.-C.iv.). ATG5 was primarily found as diffuse staining within mature Lewy bodies (A.ii., B.ii. and C.ii.) whilst LC3B labelled most Lewy bodies (A.iii., B.iii. and C.iii.). Quantitative analysis of abundance is shown in D.i. to E.iii., with each point representing the mean value per case and lines joining points from the same case to the trend of abundance across different cell types/structures within cases. Quantitative analysis of ATG5 within LBD cases identified significantly higher abundance in Lewy bodies compared to Lewy body-bearing neurons and unaffected neurons (D.i.-D.ii.). In the cingulate gyrus, ATG5 was significantly less abundant in the cytoplasm of Lewy body-bearing neurons compared to unaffected neurons from the same case (D.ii.). No enrichment of ATG5 was observed in diffuse α-synuclein aggregates (D.iii.). LC3B was enriched in Lewy bodies of the DMV compared to unaffected neurons from the same case (E.i.), whilst LC3B in the cingulate was significantly reduced in the cytoplasm of neurons with Lewy bodies compared to the cytoplasm of proximal neurons without Lewy bodies (E.ii.). LC3B was not consistently enriched in diffuse α-synuclein aggregates (E.iii.). *p<0.05, **p<0.01.

### Mitophagic cargo is found in early α-synuclein aggregates

To determine whether the incorporation of autophagic cargo into Lewy bodies represents an early or late event in the natural history of Lewy body formation, we also evaluated the abundance of the various markers in diffuse α-synuclein accumulations in DMV neurons. This analysis was based on the view that such diffuse intracellular deposits represent the first stage of α-synuclein aggregation prior to Lewy body formation [22]. This analysis demonstrated that the only marker significantly enriched in diffuse α-synuclein deposits was the mitophagy marker pUbSer65 (Figure 2 - 4).

### Lewy bodies contain aggresomal markers suggesting a regulated process of formation

To further understand the composition of Lewy bodies and to acquire a view on how and why they form, we next isolated insoluble proteins from cases for discovery proteomics analysis using data independent analysis mass spectrometry. This analysis yielded 2,816 individual proteins but in accordance with previous findings we found a substantial proteome in cases without Lewy body pathology and a subtractive method between cases and controls would likely have yielded very few hits [23]. On this basis, we elected to statistically compare proteins individually to determine those significantly enriched in the LBD cases. This analysis yielded 100 proteins significantly enriched in LBD cases that we then evaluated to determine enrichment of cellular components using Protein STRING. Cellular component enrichment analysis yielded a number of relevant components that were consistent with our analysis, with particular up-regulation of microtubules, organelles, vesicles and mitochondria (Supplementary Figure 3).

To examine functional aspects of the enriched proteome, Protein STRING analysis was performed with k-means clustering to segregate the enriched proteome into three clusters (Figure 5). This segregation combined with visual inspection of the constituent proteins and their known function suggested an immune/inflammation cluster containing caspase 1, STAT1 and NF-κB2 and a structural cluster containing tubulins and the tubulin folding complex CCT6A (Figure 5). A further cluster was identified, of which α-synuclein was a component, and contained a number of proteins related to aggresomal formation, the regulated cellular response to proteasomal impairment due to an abundance of unfolded proteins [24]. The aggresomal response consists of trafficking of unfolded proteins to the microtubule organising complex, where unfolded proteins are incorporated into a large perinuclear aggregate. In the present study, enrichment was observed of proteins implicated in trafficking misfolded proteins to the aggresome, such as HDAC6 and BAG6, proteins involved in the formation of aggresomes (UFD1) and proteasomal proteins implicated in aggresomal clearance (PSMD14 and HUWE1) [25-29] (Figure 5). Taken together, it was felt that the insoluble proteome enriched in LBD cases could be consistent with an aggresome-type response, and thus the presence of the aggresome markers HDAC pSer22 and γ-tubulin were evaluated in LBD cases. Two raters viewed Lewy bodies and diffuse α-synuclein accumulations to determine whether they were immunopositive or negative for both markers. This analysis identified that of 55 DMV Lewy bodies (16 of which were “classic” Lewy bodies), 100% contained both HDAC6 pSer22 and γ-tubulin. In contrast, of 16 diffuse α-synuclein accumulations, 31.25% contained HDAC6 pSer22 and 75% contained γ-tubulin. In cingulate gyrus, we observed 91.3% of 23 Lewy bodies were immunoreactive for HDAC6 pSer22 and 100% for γ-tubulin.

**Figure 5:**
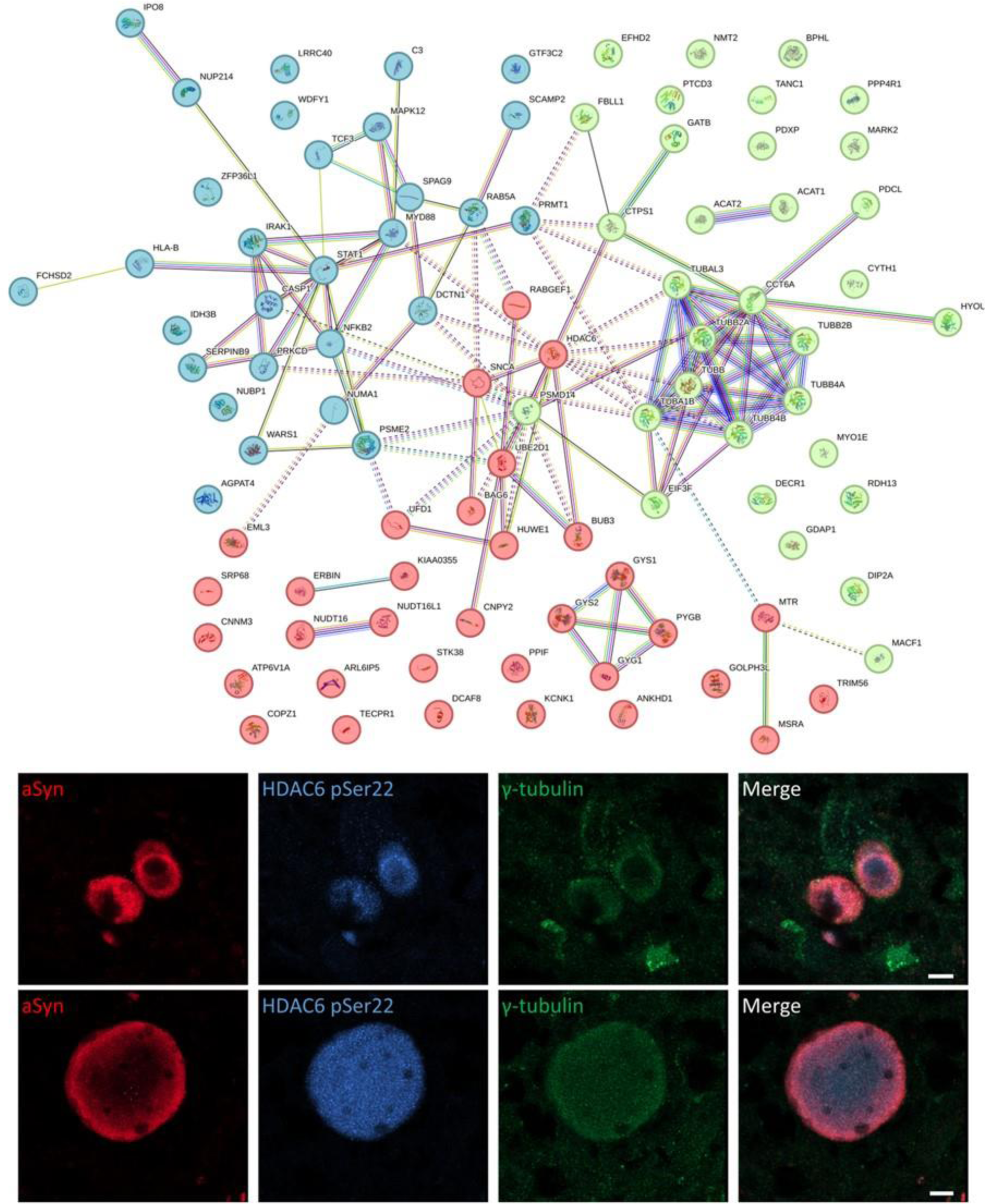
Enrichment of aggresomal markers within Lewy bodies. Proteomic analysis using STRING identified three clusters via k-means clustering, shown here in blue, green and red. The blue cluster is enriched in proteins associated with inflammation, such as NFKB2 and STAT1, whereas the green cluster contains a number of tubulins. The red cluster, which contains α-synuclein (SNCA) is enriched for aggresomal markers such as HDAC6, BAG6, UFD1, PSME2 and HUWE1. Immunofluorescent analysis of the aggresomal markers HDAC pS22 and γ-tubulin identified they are commonly found within Lewy bodies.

## DISCUSSION

The present study sought to understand why Lewy bodies occur by comparing differences between mitochondrial diseases cases with and without Lewy bodies, with a focus on impaired mitophagy as a potentially critical contributor to Lewy body formation. No differences were found in any measures of autophagy in cases with Lewy body pathology compared to cases without Lewy bodies. However, Lewy bodies were enriched for markers of autophagic vesicles and autophagic mitochondria, and this appeared to be selective for waste as no general enrichment for mitochondria was observed in Lewy bodies. Incorporation of autophagic mitochondria appeared to be an early event in Lewy body formation as mitophagy markers were up-regulated in diffuse α-synuclein aggregates thought to precede Lewy body formation. A combination of discovery proteomics and immunofluorescent analysis demonstrated that Lewy bodies are almost invariably labelled by markers of aggresomes, a mechanism of sPQC to accumulate misfolded proteins when the proteasomal system is overwhelmed. As these results suggest Lewy bodies are enriched for both cellular waste and proteins involved in trafficking proteins that cannot be degraded by conventional mechanisms, one could hypothesise that Lewy bodies are a deliberately formed structure to encapsulate and contain waste that cannot otherwise be degraded.

Autophagy is a critical cellular process to prevent both the accumulation of damaged organelles and misfolded proteins within cells, and also to salvage macromolecules to support other metabolic activities. The present study aimed to explore the hypothesis that autophagy would be deficient in individuals with Lewy body pathology, as this could suggest Lewy bodies form to encapsulate waste that cannot otherwise be degraded. This study found no consistent evidence of alterations to the abundance of any of the markers studied at the group level, perhaps suggesting limited evidence of autophagy impairment. However, it should be noted that examination of the abundance of markers of autophagic machinery is a sub-optimal approach for studying autophagy as testing abundance is not necessarily a proxy of function, particularly with regards to a dynamic approach like autophagy [30]. However, functional assays are precluded by the use of *post-mortem* human brain tissue; nevertheless, given the difficulty in generating Lewy bodies *in vitro*, which itself suggests incomplete understanding of how they form, we felt our human tissue-based approach was appropriate despite these caveats.

The observation that Lewy bodies contain autophagic mitochondria and vesicles is consistent with previous studies which have reported these as key components of Lewy bodies [12, 22]. However, our present study extends previous findings by helping elucidate further information about the identity of the vesicles and likely health of the mitochondria. Enrichment of markers of phagophores and autophagosomes within Lewy bodies suggests that vesicles carrying autophagic cargo comprise at least some of these vesicles, and the presence of organelles decorated with ubiquitin phosphorylated at serine 65 within and surrounding Lewy bodies suggests mitochondria labelled for autophagic degradation are incorporated into Lewy bodies. It was notable that there was no obvious difference in the abundance of the general mitochondrial marker VDAC1/porin within Lewy bodies compared to neuronal cytoplasm, nor reduction of mitochondrial mass in Lewy body-bearing neurons, suggesting against a general incorporation of mitochondria into Lewy bodies in a “black hole”-like manner. This does not mean mitochondria labelled by VDAC1/porin are not present in Lewy bodies, but rather than they do not show increased abundance relative to the cytoplasm, in fact they almost always showed less abundance within Lewy bodies than in the cytoplasm of the neuron in which they were located. The latter observation is not consistent with a previous *in vitro* study which showed almost complete co-localisation between the mitochondrial marker TOMM20 and aggregated α-synuclein, which was interpreted as a proposed mechanism of Lewy body-mediated toxicity in the sequestration and incorporation of healthy organelles [31]. This discrepancy between what is observed in model system and *post-mortem* human brain tissue could reflect differences between VDAC1/porin and TOMM20 antibody labelling; however, as our findings are consistent with previous observations in human brain tissue using label-free methods [12, 22], it may simply reflect different time intervals between the initiation of pathology and observation from the relative short-lived *in vitro* system and many years in human *post-mortem* tissue.

The observation that Lewy bodies were almost invariably immunoreactive for the aggresome marker HDAC6 pS22 is consistent with a previous study that reported this phenomenon in a similar number of LBD cases [32]. Aggresomes are a regulated response to misfolded proteins and proteasomal inhibition, typically in stressed cells, in which proteins are trafficked along microtubules to the microtubule organising complex by dynein motor proteins [33]. HDAC6 interacts with the dynein motor protein to transport misfolded proteins to the aggresome through the microtubule network [27], and interactions with dynein are enhanced by phosphorylation of HDAC6 at serine 22 (HDAC6 pS22) [34]. Thus, our observation that HDAC6 pS22 is enriched within Lewy bodies is consistent with the constituent components of Lewy bodies having been trafficked there via the microtubule network. Previous observations of aggresomes indicate they are enriched for filamentous proteins that form a cage reminiscent of those previously described for Lewy bodies [22, 35]. Therefore, there are striking similarities between Lewy bodies and aggresomes, suggesting the formation of Lewy bodies is a deliberate response by neurons to a stressor. Given aggresomes form in response to proteostatic stress, and the critical role ascribed to α-synuclein aggregation in LBD, one could speculate that the aggregation of α-synuclein induces proteasomal and autophagy-lysosomal dysfunction that leads to the formation of compensatory aggresomes that are rich in α-synuclein aggregates in addition to other cellular components that cannot otherwise be degraded via autophagy.

Previous studies of aggresomes have typically highlighted their enrichment for misfolded proteins and filaments that presumably provide structure to the aggresome. However, the present study, in agreement with previous observations, suggests that Lewy bodies are enriched in markers of cellular waste, including autophagosomes, autophagic mitochondria and nitrosatively modified proteins [12]. Our interpretation of this observation is that this represents an attempt by the cell to harness the inherent degradative capacity of the microtubule organising complex, around which aggresomes form, that is enriched with proteasomes, proteasomal activators and chaperones [36]. As a previous study has reported mitochondria-enriched aggresomal structures in the context of mitophagy dysfunction, the present findings could be consistent with the accumulation of autophagic cargo due to α-synuclein aggregation causing autophagy dysfunction, from which the formation of a modified aggresome termed a Lewy body is a compensatory process to encapsulate autophagic cargo [37]. An alternative explanation is that α-synuclein aggregates are translocated to aggresomes, and that the presence of autophagic markers within Lewy bodies as presently reported is due to aggregates being trafficked within autophagosomes to the aggresome, as has been reported for other misfolded proteins [35]. Furthermore, the presence of autophagic mitochondria could be due to the presence of α-synuclein aggregates in damaged mitochondria, as has also been reported previously [38].

However, whether the primary substrate of encapsulation into the Lewy body aggresome is autophagic cargo, α-synuclein aggregates, or both, the engagement of aggresomal machinery suggests the formation of these structures is a deliberate process rather than the passive end result of α-synuclein aggregation. Furthermore, as aggresomal sequestration of toxic misfolded proteins has previously been reported to be protective against a range of protein aggregates, including α-synuclein, this process could be adaptive and compensatory rather than harmful [24, 39-42]. These observations have important implications, as they challenge the central pathogenic role ascribed to Lewy bodies in LBD that is the basis of many neuropathological studies and all neuropathological staging systems for Lewy body pathology.

We acknowledge a number of limitations in the present study. Firstly, the cohort size is small, though this reflects the relative rarity of mitochondrial disease brains, particularly those of relatively advanced age and thus likely to manifest Lewy bodies. Furthermore, the analysis performed was detailed, focusing on individual neurons, and study power was enhanced by comparisons within individuals. Nevertheless, the extent to which the present findings can be generalised beyond mitochondrial disease cases with Lewy bodies to idiopathic LBD cases is not clear. Our previous study suggested that Lewy bodies in mitochondrial disease conform to the same topographical progression scheme as LBD and are immunoreactive for post-translational markers of α-synuclein observed in Lewy bodies in LBD, suggesting Lewy bodies in mitochondrial disease are the same or very similar to those in LBD [15]. To attempt to overcome this potential limitation, we included two PDD cases and one incidental LBD case, which all showed very similar findings to mitochondrial disease cases with Lewy bodies; however, only replication in a large cohort of LBD cases will confirm if the present findings are observed across LBD cases.

In conclusion, neurons from mitochondrial disease cases with Lewy bodies do not show abnormalities in any autophagy markers compared to neurons from individuals with mitochondrial disease without Lewy bodies. However, neurons with Lewy bodies show striking differences from unaffected neurons, including incorporation of a wide array of markers of autophagic cargo and aggresomes which, taken together, could suggest Lewy bodies are modified aggresomes. As aggresomes have been demonstrated to be a highly regulated and protective mechanism, one could speculate that Lewy bodies are purposefully formed to encapsulate misfolded α-synuclein and other cellular waste, thus reflecting an adaptive and perhaps even protective process.

## ACKNOWLEDGEMENTS

This study was funded by the Alzheimer’s Research UK Senior Fellowship awarded to DE (ARUK-SRF2022A-006), with support from the Wellcome Centre for Mitochondrial Research at Newcastle University (203105/Z/16/Z) and the UK NHS Specialist Commissioners which funds the “Rare Mitochondrial Disorders of Adults and Children” Diagnostic Service in Newcastle upon Tyne.

## SUPPLEMENTARY DATA

**Supplementary Figure 1:**
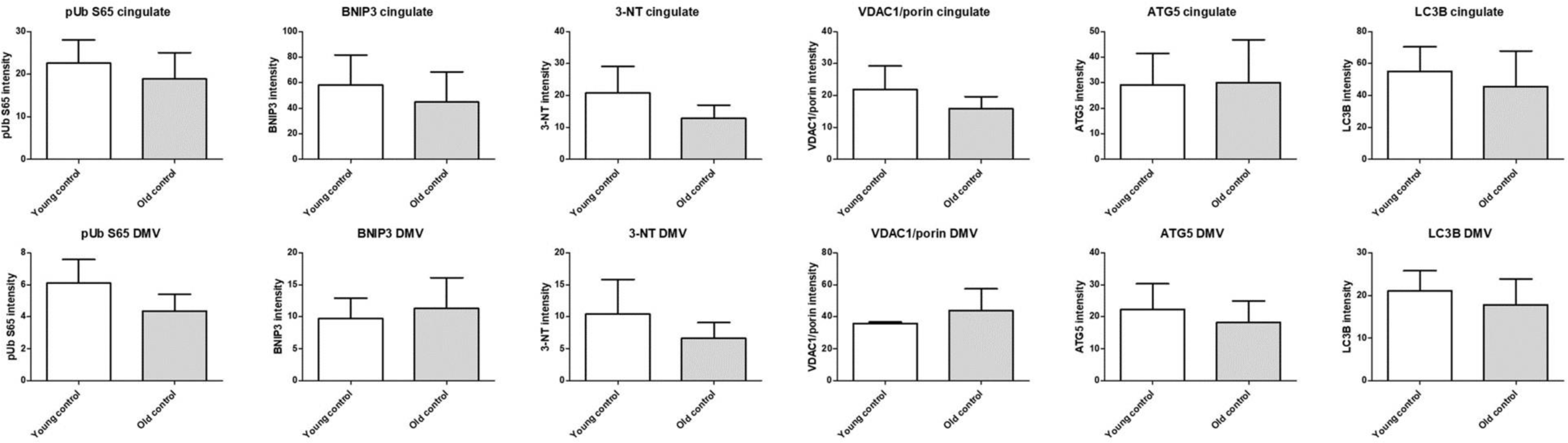
Bar charts demonstrating no difference in neuronal abundance of all markers between young and old control cases.

**Supplementary Figure 2:**
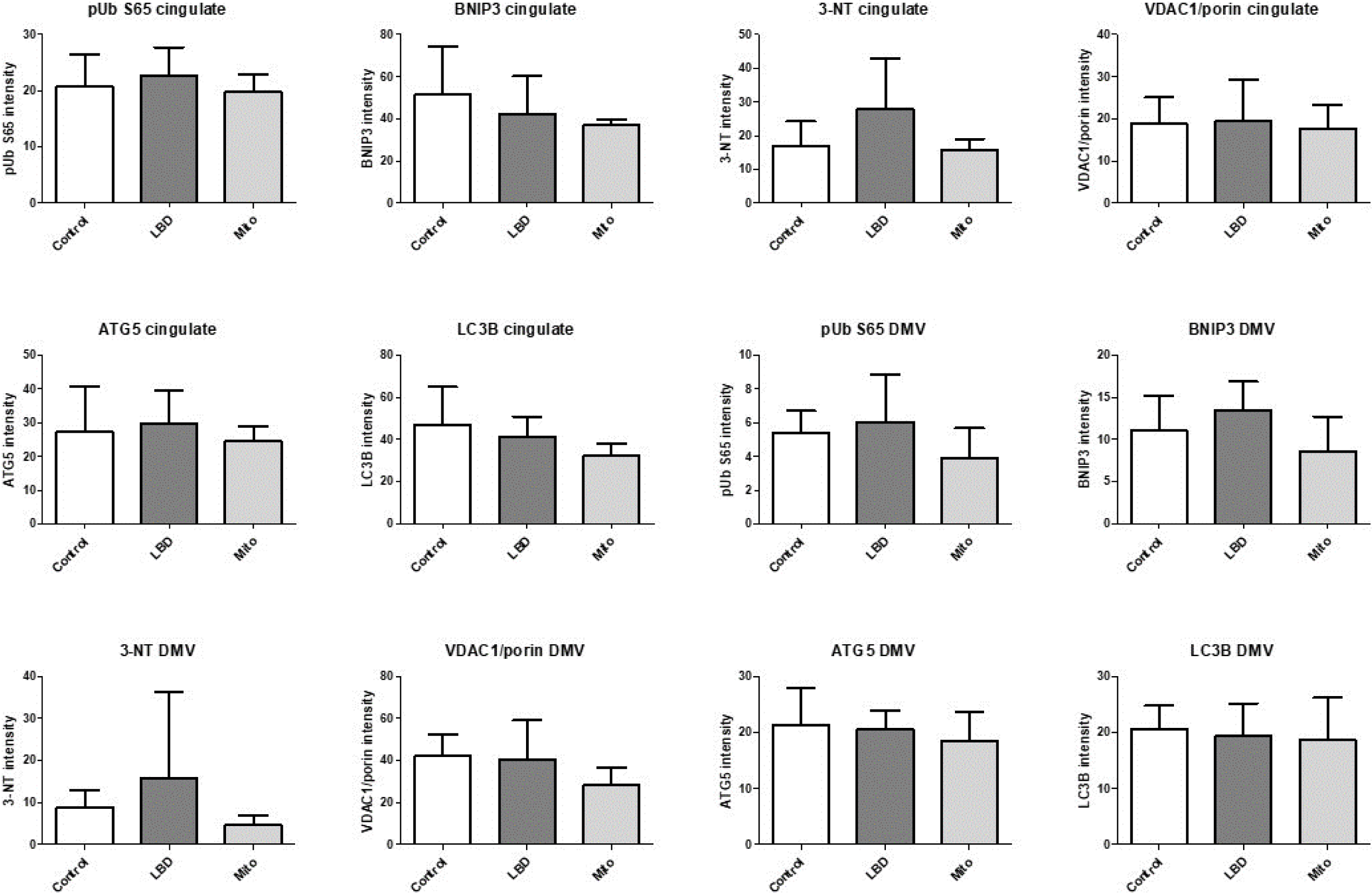
Bar charts showing abundance levels of different marks at the group level. None were statistically significant.

## Notes

### Competing Interest Statement

V.I.K. is a Scientific Advisor for Longaevus Technologies. All other authors declare they have no competing interests.

