## Supplementary Figure 3 for "Autophagic cargo in Lewy bodies: are Lewy bodies a compartment for spatial protein quality control?"

| LBD mean | Control mean | Normalised difference | Gene | Protein |
|---|---|---|---|---|
| 3.55688 | 0.183222167 | 3.373657833 | SNCA | Alpha-synuclein |
| 2.3806 | -0.9548405 | 3.3354405 | FBLL1 | rRNA/tRNA 2'-O-methyltransferase |
| -0.354339 | -3.368265 | 3.01392565 | GARRE1 | Granule associated Rac and RHOG |
| 6.436604 | 3.61631 | 2.820294 | GYG1 | Glycogenin-1 |
| 0.352368 | -2.454113667 | 2.806481667 | ACAT2 | Acetyl-CoA acetyltransferase, cyto |
| 0.5430117 | -1.983573333 | 2.526585 | PPIE;PPIF | Peptidyl-prolyl cis-trans isomerase |
| 0.996499 | -1.43191775 | 2.42841675 | GYS2 | Glycogen [starch] synthase, liver |
| -0.623329 | -3.00363 | 2.3803015 | SPAG9 | C-Jun-amino-terminal kinase-inter |
| -1.60165 | -3.941755 | 2.340105 | AGPAT4 | 1-acyl-sn-glycerol-3-phosphate acy |
| -2.25556 | -4.430535 | 2.174975 | SCAMP2 | Secretory carrier-associated meml |
| -0.325579 | -2.477671333 | 2.152092 | SRP68 | Signal recognition particle subunit |
| -1.890337 | -4.03418 | 2.143843333 | STK38 | Serine/threonine-protein kinase 3 |
| -2.077236 | -4.20874 | 2.13150376 | WARS1 | Tryptophan--tRNA ligase, cytoplas |
| -1.267409 | -3.393416667 | 2.126007417 | COPZ1 | Coatomer subunit zeta-1 |
| 0.4550463 | -1.6400372 | 2.09508345 | HLA-B | HLA class I histocompatibility antig |
| -1.150995 | -3.22462 | 2.073625 | PPP4R1 | Serine/threonine-protein phospha |
| -0.83302 | -2.78594 | 1.95292 | MARK2 | Non-specific serine/threonine prot |
| -2.174357 | -4.11244 | 1.938083333 | ARL6IP5 | PRA1 family protein 3 |
| -1.217425 | -3.051408 | 1.833982667 | RABGEF1 | Rab5 GDP/GTP exchange factor |
| -0.077963 | -1.881762694 | 1.803799694 | PRMT1 | Protein arginine N-methyltransfera |
| -2.75356 | -4.47323 | 1.71967 | GOLPH3L | Golgi phosphoprotein 3-like |
| 3.086428 | 1.373226333 | 1.713201667 | GYS1 | Glycogen [starch] synthase, muscl |
| -1.85423 | -3.539485 | 1.685255 | CNNM3 | Metal transporter CNNM3 |
| 1.2825994 | -0.376998833 | 1.659598233 | ERBIN | Erbin |
| -1.439916 | -3.089715 | 1.649798667 | NUBP1 | Cytosolic Fe-S cluster assembly fac |
| -2.482563 | -4.075695 | 1.5931325 | IPO8 | Importin-8 |
| -0.403707 | -1.98422 | 1.580513267 | PSME2 | Proteasome activator complex sub |
| -2.34306 | -3.8885725 | 1.5455125 | PTCD3 | Pentatricopeptide repeat domain- |
| -0.930161 | -2.46842 | 1.5382594 | NUDT16 | U8 snoRNA-decapping enzyme |
| 2.799122 | 1.267630167 | 1.531491833 | LRRC40 | Leucine-rich repeat-containing pro |
| -1.55299 | -3.043034 | 1.490044 | GATB | Glutamyl-tRNA(Gln) amidotransfer |
| -0.183019 | -1.6688714 | 1.4858525 | MAPK12 | Mitogen-activated protein kinase |
| -0.229502 | -1.696489667 | 1.466987333 | BPHL | Valacyclovir hydrolase |
| -0.539433 | -1.9537534 | 1.414320067 | ANKHD1;AI | Ankyrin repeat domain-containing |
| -0.772815 | -2.184725 | 1.41191 | TANC1 | Protein TANC1 |
| 0.345366 | -1.0257945 | 1.3711605 | TCF3 | Transcription factor E2-alpha (Frag |
| 1.973802 | 0.613326167 | 1.360475833 | TUBAL3 | Tubulin alpha chain-like 3 |
| -1.61128 | -2.968864 | 1.357584 | DCAF8 | DDB1- and CUL4-associated factor |
| 0.7687186 | -0.568912367 | 1.337630942 | WDFY1 | WD repeat and FYVE domain-cont |
| -1.973565 | -3.301315 | 1.32775 | SERPINB9 | Serpin B9 |
| -0.449059 | -1.763908333 | 1.314849083 | CASP1 | Caspase-1 |
| -0.474931 | -1.772042 | 1.2971108 | ZFP36L1 | mRNA decay activator protein ZFP |
| -0.18325 | -1.464091667 | 1.280841917 | EIF3F | Eukaryotic translation initiation fac |
| 0.7308213 | -0.546509833 | 1.277331113 | TUBB4A | Tubulin beta-4A chain (Fragment) |
| -0.086557 | -1.3628645 | 1.276307425 | EML3 | Echinoderm microtubule-associate |
| -1.1827 | -2.416718333 | 1.234018583 | CNPY2 | Protein canopy homolog 2 |
| -0.834565 | -2.050706667 | 1.216142167 | NUMA1 | Nuclear mitotic apparatus protein |
| -0.715946 | -1.893726667 | 1.177780267 | NUP214 | Nuclear pore complex protein Nup |
| -1.476664 | -2.644388 | 1.1677238 | DIP2A | Disco-interacting protein 2 homolo |

| | | | | |
|---|---|---|---|---|
| -0.19527 | -1.3158446 | 1.120575 | GTF3C2 | General transcription factor 3C po |
| -2.130545 | -3.234933333 | 1.104388333 | NMT2 | Glycylpeptide N-tetradecanoyltrar |
| -1.283492 | -2.386402 | 1.1029102 | PSMD14 | 26S proteasome non-ATPase regul |
| -1.368726 | -2.456416 | 1.08769 | DECR1 | 2,4-dienoyl-CoA reductase [(3E)-er |
| -1.594355 | -2.642036667 | 1.047681667 | DCTN1 | Dynactin subunit 1 |
| 1.9146073 | 0.9010228 | 1.01358445 | PPIF | Peptidyl-prolyl cis-trans isomerase |
| -0.34753 | -1.3524875 | 1.00495752 | MSRA | Mitochondrial peptide methionine |
| -0.948122 | -1.952954 | 1.0048325 | UFD1 | Ubiquitin recognition factor in ER-a |
| -0.68407 | -1.688536667 | 1.004466667 | PDCL | Phosducin-like protein |
| 1.2179902 | 0.2451351 | 0.9728551 | ACAT1 | Acetyl-CoA acetyltransferase, mitc |
| -1.34046 | -2.27272 | 0.93226 | MTR | Methionine synthase |
| -0.635623 | -1.566507833 | 0.930884833 | MACF1 | Microtubule-actin cross-linking fac |
| -1.803425 | -2.71407 | 0.910645 | UBE2D1;UE | Ubiquitin-conjugating enzyme E2 D |
| -1.731113 | -2.63783 | 0.9067175 | CYTH1 | Cytohesin-1 |
| 1.64877 | 0.777498833 | 0.871271167 | GDAP1 | Ganglioside-induced differentiatio |
| -0.399109 | -1.2532666 | 0.8541576 | BUB3 | Mitotic checkpoint protein BUB3 |
| -1.9301 | -2.7814 | 0.8513 | MYD88 | Myeloid differentiation primary re |
| 2.261114 | 1.487524667 | 0.773589333 | IDH3B | Isocitrate dehydrogenase [NAD] su |
| -0.555705 | -1.297600333 | 0.741895083 | HDAC6 | Histone deacetylase 6 |
| -0.765566 | -1.500695 | 0.7351295 | NFKB2 | Nuclear factor NF-kappa-B p100 su |
| 1.3654466 | 0.6336895 | 0.7317571 | C3 | Complement C3 |
| -0.081497 | -0.809558577 | 0.728061337 | EFHD2 | EF-hand domain-containing protei |
| 0.5619542 | -0.1660735 | 0.7280277 | PDXP | Chronophin |
| -1.021495 | -1.678035 | 0.65654025 | MYO1E;MY | Unconventional myosin-If |
| 7.894914 | 7.24655 | 0.648364 | TUBB;TUBE | Tubulin beta chain |
| 5.660904 | 5.013836667 | 0.647067333 | TUBA1C;TU | Tubulin alpha-1C chain |
| -0.584954 | -1.2266828 | 0.6417288 | TRIM56 | E3 ubiquitin-protein ligase TRIM56 |
| 0.472762 | -0.1567375 | 0.62949952 | CTPS1 | CTP synthase 1 |
| -0.127823 | -0.750294333 | 0.622471533 | PRKCD | Protein kinase C delta type |
| -0.917002 | -1.538075 | 0.6210728 | NUDT16L1 | Tudor-interacting repair regulator |
| 0.815912 | 0.1972121 | 0.6186999 | RDH13 | Retinol dehydrogenase 13 |
| 2.03168 | 1.413297 | 0.618383 | CCT6A | T-complex protein 1 subunit zeta |
| 9.90993 | 9.323495 | 0.586435 | TUBB2A;TU | Tubulin beta-2A chain |
| -0.955133 | -1.5334675 | 0.5783349 | TECPR1 | Tectonin beta-propeller repeat-co |
| 8.954208 | 8.38103 | 0.573178 | TUBB2A | Tubulin beta-2A chain |
| 1.2837166 | 0.714600667 | 0.569115933 | HUWE1 | E3 ubiquitin-protein ligase HUWE1 |
| 11.9617 | 11.41535 | 0.54635 | TUBA1B;TU | Tubulin alpha-1B chain |
| 10.26244 | 9.776176667 | 0.486263333 | TUBB;TUBE | Tubulin beta-4A chain |
| 10.089274 | 9.60656 | 0.482714 | TUBA1B;TU | Tubulin alpha-1B chain |
| 3.332396 | 2.8588 | 0.473596 | ATP6V1A | V-type proton ATPase catalytic sub |
| -0.690425 | -1.160532333 | 0.470107333 | IRAK1 | Interleukin-1 receptor-associated k |
| 3.52162 | 3.052436667 | 0.469183333 | KCNK1 | Potassium channel subfamily K me |
| 9.384 | 8.9214 | 0.4626 | TUBB | Tubulin beta chain |
| 7.199966 | 6.751935 | 0.448031 | TUBA1B;TU | Tubulin alpha-1B chain |
| -0.994171 | -1.431643333 | 0.437472583 | FCHSD2 | F-BAR and double SH3 domains pr |
| 1.377702 | 0.941147833 | 0.436554167 | RAB5A | Ras-related protein Rab-5A |
| -1.283248 | -1.7007825 | 0.4175343 | HYOU1 | Hypoxia up-regulated protein 1 |
| 2.412498 | 2.0048 | 0.407698 | STAT1 | Signal transducer and activator of |
| 8.139086 | 7.73338 | 0.405706 | TUBB4B | Tubulin beta-4B chain |
| 3.958652 | 3.592956667 | 0.365695333 | PYGB | Glycogen phosphorylase, brain for |

| | | | | |
|---|---|---|---|---|
| 0.4246114 | 0.133697733 | 0.290913667 | BAG6 | Large proline-rich protein BAG6 |

| Cellular component | Strength |
|---|---|
| Microtubule bundle | 2.03 |
| Mitotic spindle | 0.85 |
| Microtubule | 0.75 |
| Polymeric cytoskeletal fibre | 0.57 |
| Supramolecular fibre | 0.48 |
| Supramolecular complex | 0.43 |
| Mitochondrion | 0.38 |
| Extracellular exosome | 0.37 |
| Cytosol | 0.34 |
| Cytoplasmic vesicle | 0.32 |

| FDR |
| --- |
| 0.0282 |
| 0.025 |
| 0.00055 |
| 0.0126 |
| 0.025 |
| 0.0218 |
| 0.0252 |
| 0.0147 |
| 1.26E-07 |
| 0.0313 |
